# The importance of stereochemistry in the disorder-order continuum of protein-protein interactions

**DOI:** 10.1101/2024.02.23.581681

**Authors:** Estella A. Newcombe, Amanda D. Due, Andrea Sottini, Catarina B. Fernandes, Lasse Staby, Elise Delaforge, Christian R. O. Bartling, Inna Brakti, Katrine Bugge, Benjamin Schuler, Karen Skriver, Johan G. Olsen, Birthe B. Kragelund

## Abstract

Intrinsically disordered proteins can bind *via* the formation of highly disordered protein complexes without the formation of 3D-structure. Most naturally occurring proteins are “left-handed” or levorotatory (L), made up only of L-amino acids, imprinting molecular structure and communication with stereochemistry. In contrast, their mirror image “right-handed” or dextrorotatory (D) amino acids are rare in Nature. Whether disordered protein complexes are truly independent of 3D-topology and thus of chiral constraints is not clear. To test the chiral constraints of disordered protein-protein interactions, a set of interacting protein pairs covering the disorder-order continuum was chosen as representative examples. By observing both the natural ligands and their stereochemical mirror images in free and bound states, we discovered that chirality was inconsequential in a fully disordered complex. However, if the interaction relied on the ligand undergoing coupled folding and binding, correct stereochemistry was essential. Between these extremes, binding could be observed for the D-ligand with a strength that correlated with the amount of disorder in the final complex. These findings have important implications for our understanding of protein-protein interactions, the molecular processes leading to complex formation, the use of D-peptides in drug discovery, and the chemistry of protein evolution of the first living entities on Earth.

## INTRODUCTION

The stereochemistry of amino acids, and therefore of proteins, is biological canon. The chirality of the C^*α*^-atom means that the mirror images (enantiomers) of amino acids cannot be superimposed; they have a “handedness”. Amino acids in Nature are predominantly “left-handed” or levorotatory (L), whereas their enantiomers are “right-handed” or dextrorotatory (D) (**Fig. 1A**) - so named because of the way they affect circularly polarized light^1^. Thus, L- and D-amino acids, and hence L- and D-amino acid-based proteins (L- and D-proteins), are mirror images (**Fig. 1B**). The preference for L-proteins is so strong that we may generally say that L-proteins make up the molecular structure and machinery of Nature. However, D-amino acids do exist, and Nature exploits these typically in signaling as free amino acids, or in defense systems as parts of smaller peptides or peptidoglycans, e.g., in the bacterial cell wall^2,3^, as neurotransmitters^4^, toxins and venoms^5^, and antibiotics^6^ (for a review see^7^).

**Figure 1.**
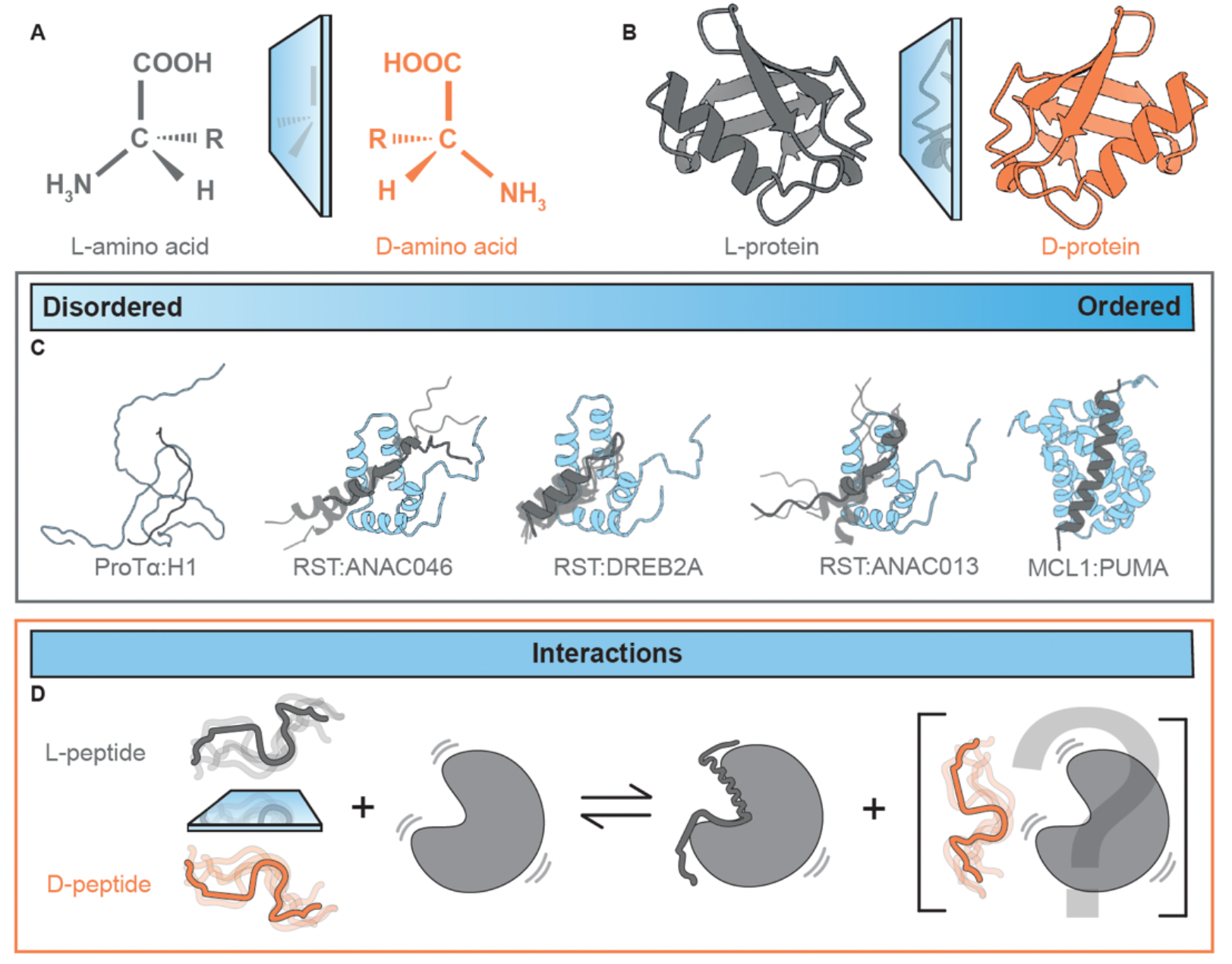
Chirality in protein-protein interactions. **A** L- and D-amino acids are mirror images, as are **B** L- and D-proteins. **C** Model systems covering a continuum of disordered (ProTα:H1)^15,16^ to ordered (MCL1:PUMA)^17^ protein complexes, and three intermediate interactions with RST: ANAC046, DREB2A^18^ and ANAC013. **D** L-protein pairs will interact, but the features that might allow L- and D-proteins to interact are unclear.

Proteins are key to the activity of biological systems, functioning via interactions with one or several binding partners. It is widely accepted that the D-enantiomer of a protein would be unable to bind a partner L-protein. However, in a pharmaceutical context, it would be desirable to overcome this lack of binding due to the metabolic stability of D-proteins in biological systems, where they are not recognized by natural metabolic processes^8^. Thus, D-amino acid-based peptides have been explored as constituents of peptide drugs and synthetic D-proteins as scaffold for screening L-peptides in mirror-image phage-diplays^9^. In the “retro-inverso” strategy, D-amino acid-based peptides mimic the L-peptide enantiomer when producing the D-amino acid sequence in reverse^10^. This strategy relies on the D-peptide forming the same secondary structure as the L-peptide, enabling interaction with its L-protein binding partners. Examples can be found in the treatment of diabetes^11,12^, breast cancer^13^, and inflammation^14^.

The last 25 years have uncovered the functional relevance of intrinsically disordered proteins and protein regions (collectively here IDPs) existing in dynamic ensembles of interconverting conformations^19^. Although structural disorder can remain in complexes and plays important functional roles there, it remains unclear whether IDPs are also confined to chiral structural constraints^20–22^. The continuum of complexes formed by IDPs range from folded, induced-fit interactions to fuzzy, or fully disordered, complexes with structural heterogeneity^15,23^. Not much is known about the atomic structure of heterogeneous complexes, and less so of the fully disordered ones where the ligand at the extreme can be comparably dynamic in the free and the bound states^15,24^. This raises the fundamental question of whether these complexes are truly independent of 3D-topology and thus independent of the chiral constraints of folded complexes, or whether there are configurational constraints, perhaps too subtle to be experimentally resolved.

To answer this question, we selected an assortment of interacting protein pairs in which the ligand is disordered in its unbound state, and either stays disordered in the complex or adopts different degrees of structure upon binding (**Fig. 1C**). Peptide ligands were synthesized using D-amino acids and compared to their L-peptide enantiomers using a range of biophysical and structural methodologies (**Fig. 1D**). We found that sensitivity to chirality in binding correlates with the degree of folding in the complex, with disordered protein complexes forming regardless of the “handedness” of the ligand.

## RESULTS

To test whether disordered protein interactions could persist regardless of chirality, we initially focused on the prothymosin-α (ProTα) interaction with histone 1.0 (H1), which has been shown to be a high-affinity, disordered interaction^15,16^ (**Fig. 1C**). We used a 21-residue peptide from the C-terminal tail of H1_155-175_ (**Fig. 2A**), containing a high charge density with a fraction of charged residues (FCR) of 0.52 (**Table S1**), and procured both an L- and D-enantiomer (L-H1_155-175_ and D-H1_155-175_, respectively). Far-UV circular dichroism (CD) confirmed the two peptides as mirror images (**Fig. 2B**), and nuclear magnetic resonance (NMR) spectroscopy analyses showed identical chemical shifts (**Fig. S1A**). The CD spectra also showed that the peptides were disordered, as expected. We next used NMR to measure the chemical shift perturbations (CSPs) of ProTα caused by each enantiomer upon their addition. In this case, we found that L- and D-H1_155-175_ produced similar CSPs in ProTα (**Fig. 2C**), which we quantified by calculating the difference between the CSPs (ΔCSPs) at equimolar concentrations of each enantiomer of the H1_155-175_ peptide (**Fig. 2D**).We probed the affinity (*K*_*d*_) and thermodynamic properties using isothermal titration calorimetry (ITC), finding the same values for the enantiomers in terms of *K*_*d*_ (**Fig. 2E, Table S2**). We observed that the changes in binding enthalpy (-TΔ*S*) and entropy (Δ*H*) were similar for L- and D-H1_155-175_ (**Table S2**), and that the *K*_*d*_ was within the low micro-molar range for both enantiomers. ProTα:H1 interact with nano-molar to picomolar affinity^15,16^, but we observed micro-molar affinity with the peptides, mostly because of the lower total charge of the H1_155-175_ fragment (ProTα -43; L/D-H1_155-175_ +11; full-length H1.0 +53). Thus, charge imbalance likely underlies this difference in affinity when compared to full-length H1. We also obtained binding affinities using single-molecule Förster resonance energy transfer (smFRET) spectroscopy, labeling ProTα with donor and acceptor fluorophores (**Fig. 2F**). The agreement between the affinities obtained by ITC and smFRET using very different ProTα concentrations suggests that the complex is predominantly of 1:1 stoichiometry (**Fig. 2**). Furthermore, analogous to the NMR CSPs, the smFRET data showed that the changes in transfer efficiencies on binding were very similar for L- and D-H1_155-175_, indicating that the conformational ensemble of ProTα bound to an L- or D-H1_155-175_ is highly similar, again highlighting that there is no significant difference between the interactions of ProTα with the L- or D-enantiomers of H1_155-175_.

**Figure 2.**
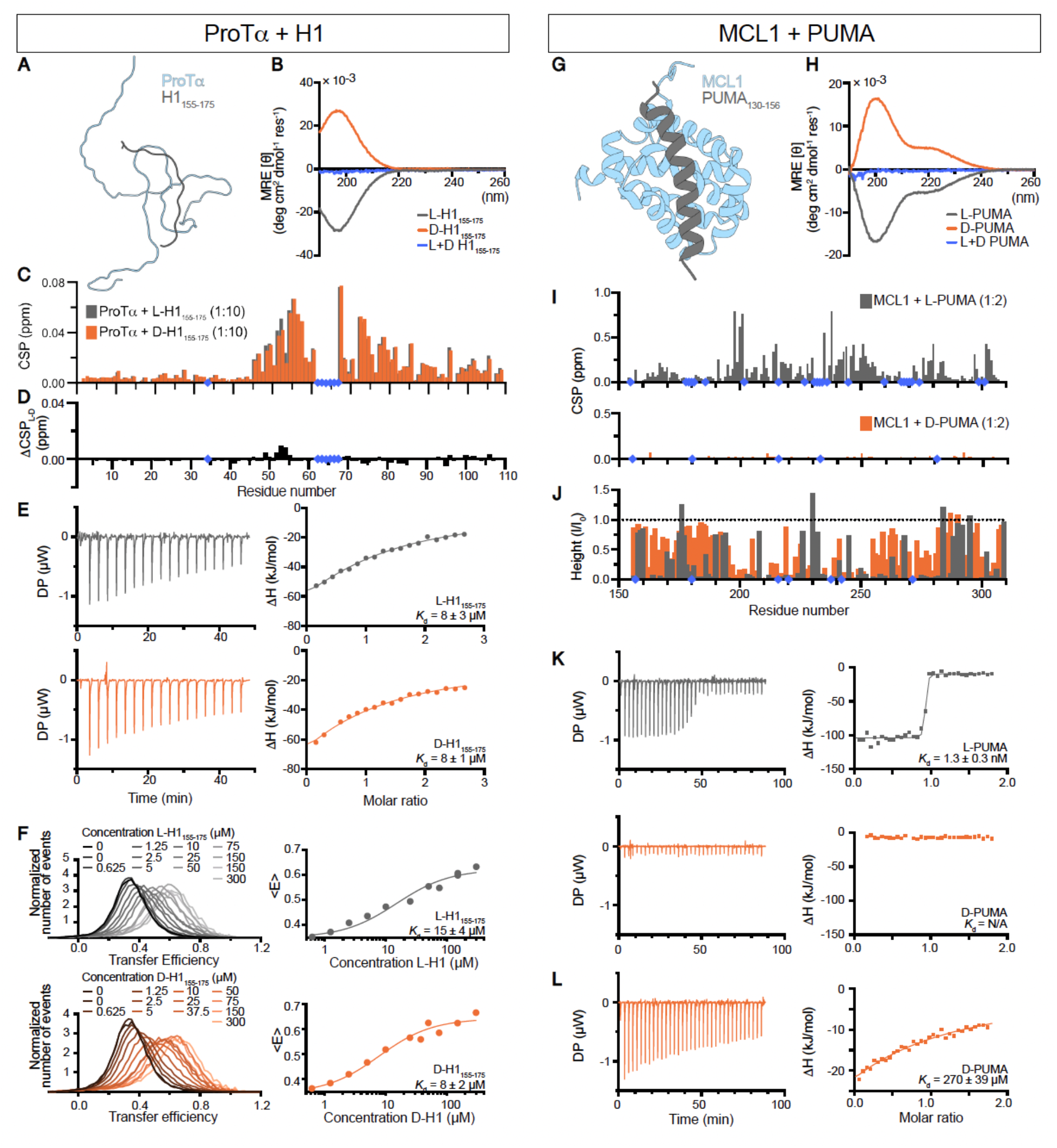
Effects of chirality on protein interactions. **A** ProTα (light grey) and the H1_155-175_ peptide (dark grey) remain disordered during interaction. **B** Far-UV CD spectra of L- and D-H1_155-175_ peptides as well as their sum (blue). **C** Chemical shift perturbations (CSPs) caused by the addition of either L- or D-H1 to ProTα, with the difference between L- and D-H1_155-175_ induced CSPs (*ΔCSP*_*L-D*_) (**D**). **E** ITC of ProTα’s interactions with L- and D-H1_155-175_ with the left side showing the raw ITC thermogram and the right side showing the fitted one-site binding isotherms. **F** Single molecule FRET of L- or D-H1_155-175_ with ProTα, fitting <*E*> to obtain *K*_*d*_. **G** MCL1 (light grey) is folded and interacts with the disordered PUMA peptide (dark grey), forming an α-helix upon interaction *via* induced fit. **H** Far-UV CD spectra of L- and D-PUMA peptides, and their sum (blue). **I** CSPs induced by the interaction of MCL1 with L- and D-PUMA at a ratio of 1:2. **J** Changes in NMR peak intensities upon addition of L- or D-PUMA to MCL1. **K** ITC performed under the same conditions for both L- and D-PUMA. **L** ITC performed using higher concentrations of both MCL1 and D-PUMA. In all panels, L-peptides are represented in grey, and D-peptides in orange. Blue diamonds indicate missing assignments, assigned residues that could not be tracked or prolines.

We next probed the interaction between induced myeloid leukemia cell differentiation protein (MCL1) and p53 upregulated modulator of apoptosis (PUMA, also known as Bcl-2-binding component 3) L- and D-peptides (**Table S1**). This nano-molar affinity complex has previously been characterized as a folding-upon-binding induced fit interaction, leading to the folding of disordered PUMA_130-156_into an α-helix within the complex^17^ (**Fig. 2G**). We therefore hypothesized that it would be unlikely for the D-enantiomer of PUMA to bind MCL1. As PUMA tends to form dimers, we used the monomeric M144I-variant^25^. As judged by far-UV CD and NMR analyses, this variant was disordered, and enantiomers produced mirror image CD spectra and identical chemical shifts (**Fig. 2H, Fig. S1**). When we added L- or D-PUMA to MCL1, only L-PUMA caused substantial CSPs (**Fig. 2I**). The interaction between MCL1 and L-PUMA was in slow exchange on the NMR timescale, whereby peaks disappear and reappear at different chemical shifts with intensities proportionally to how much protein is in each state. Surprisingly, the changes in the intensity of peaks originating from free MCL1 occur for both L- and D-PUMA, with less intensity loss observed for D-PUMA (**Fig. 1J**). Although the peaks lose intensity upon the addition of D-PUMA to MCL1, we did not see reappearance of peaks at new positions. This might indicate that D-PUMA can form an encounter complex with MCL1 but cannot undergo the folding required for an induced fit interaction. Finally, we performed ITC at the same concentrations for both L- and D-PUMA, finding that L-PUMA bound with nano-molar affinity, while D-PUMA appeared not to bind (**Fig. 2K, Table S2**). By increasing the concentrations of both MCL1 and D-PUMA considerably, we were able to observe a *K*_d_ in the high micro-molar to low milli-molar range. All thermodynamic properties (**Table S2**) were significantly different for L- and D-PUMA, which further suggest that there is a significant difference between L- and D-PUMA in their ability to interact with MCL1.

Overall, using L- and D-peptides and comparing interactions at the extremes of the disorder-order continuum show that chirality matters little in a fully disordered interaction, but is compulsory for an interaction that relies on structure.

Having probed the extremes of the disorder continuum, it was important to understand how intermediate systems behave and respond to chirality. As intermediate systems, we used the RCD1-SRO-TAF4 (RST) domain from Radical-Induced Cell Death1 (RCD1), which interacts with various transcription factors that form different degrees of structure in their RST-bound states^26^ (**Fig. 1C**). We characterized RST interactions with the transcription factor peptides ANAC046_319-338_ (**Fig. 3A inset**), DREB2A_255-272_ (**Fig. 3B inset**) and ANAC013_254-274_ (**Fig. 3C inset**; **Table S1**). Previous far-UV CD analyses suggested induced structure for the DREB2A complex with RCD1-RST, but not for the two other ligands^26^. While a data-driven model exists for the RST:DREB2A complex (**Fig. 3B inset**)^18^, the structures formed by ANAC046 and ANAC013 in their complexes with RST are unknown. Thus, for visual representation, we generated a structural prediction for these interactions using Colabfold (**Fig. 3A, C insets, respectively**)^27^. We first confirmed via far-UV CD and NMR that the L- and D-peptides were disordered, and true enantiomers (**Fig. 3A, B, C, Fig. S1**). To determine the affinity of the interactions of L- or D-peptides with RST, we used ITC (**Fig. 3D, E, F; Table S2**). We observed very different thermodynamic profiles for the three L-peptides, with more favorable enthalpy (Δ*H*°) for DREB2A and ANAC013 than for ANAC046, while the opposite was the case for the entropy (-TΔ*S*°)^28^. The thermodynamic data, in addition to the CSPs, suggest more structuring in RST-complexes with DREB2A and ANAC013 than with ANAC046. Comparing the effect of stereochemistry, we observed larger differences in *K*_d_ as the interactions became more structured, i.e., the difference between L- and D-ANAC046 was only 15-fold (**Fig. 3D**), between L- and D-DREB2A 72-fold (**Fig. 3E**), and between L- and D-ANAC013 500-fold (**Fig. 3F**). The trend indicates that the amount of structure required for binding reduces the propensity of the D-enantiomer for interacting with RST. This interpretation was further confirmed by calculating the differences in CSPs (ΔCSPs_(L-D)_) for ANAC046 (**Fig. 3H**), DREB2A (**Fig. 3I**) and ANAC013 (**Fig. 3J**). The ΔCSPs_(L-D)_ were substantial for ANAC013 and minimal for ANAC046, indicating that ANAC046 is relatively disordered in complex with RST, while ANAC013 is more structured, and therefore less likely to interact with RST as a D-enantiomer. The DREB2A ΔCSPs_(L-D)_ lie between those of ANAC046 and ANAC013, consistent with the differences in binding affinity. Finally, as our data suggested formation of an encounter complex between D-PUMA and MCL-1 but no subsequent folding, we addressed whether any of the RST ligands showed similar behavior. We extracted *K*_d_ and *k*_*off*_ after fitting NMR titration data to a two-state model using NMR 2D lineshape analysis (**Fig. S2**), finding only minor effects of stereochemistry on the transition state energies (between -2 and 1 kJ mol^-1^ (ΔΔG_unbound-‡,(D-L)_, **Table S3**)). This observation highlights that the major effect of stereochemistry occurs after the encounter complex has formed, in agreement with the observation of encounters made with D-PUMA in its interaction with MCL-1. Overall, we have systematically shown that in the interactions between L- and D-proteins the sensitivity to stereochemistry depends on the extent of disorder in the complex.

**Figure 3.**
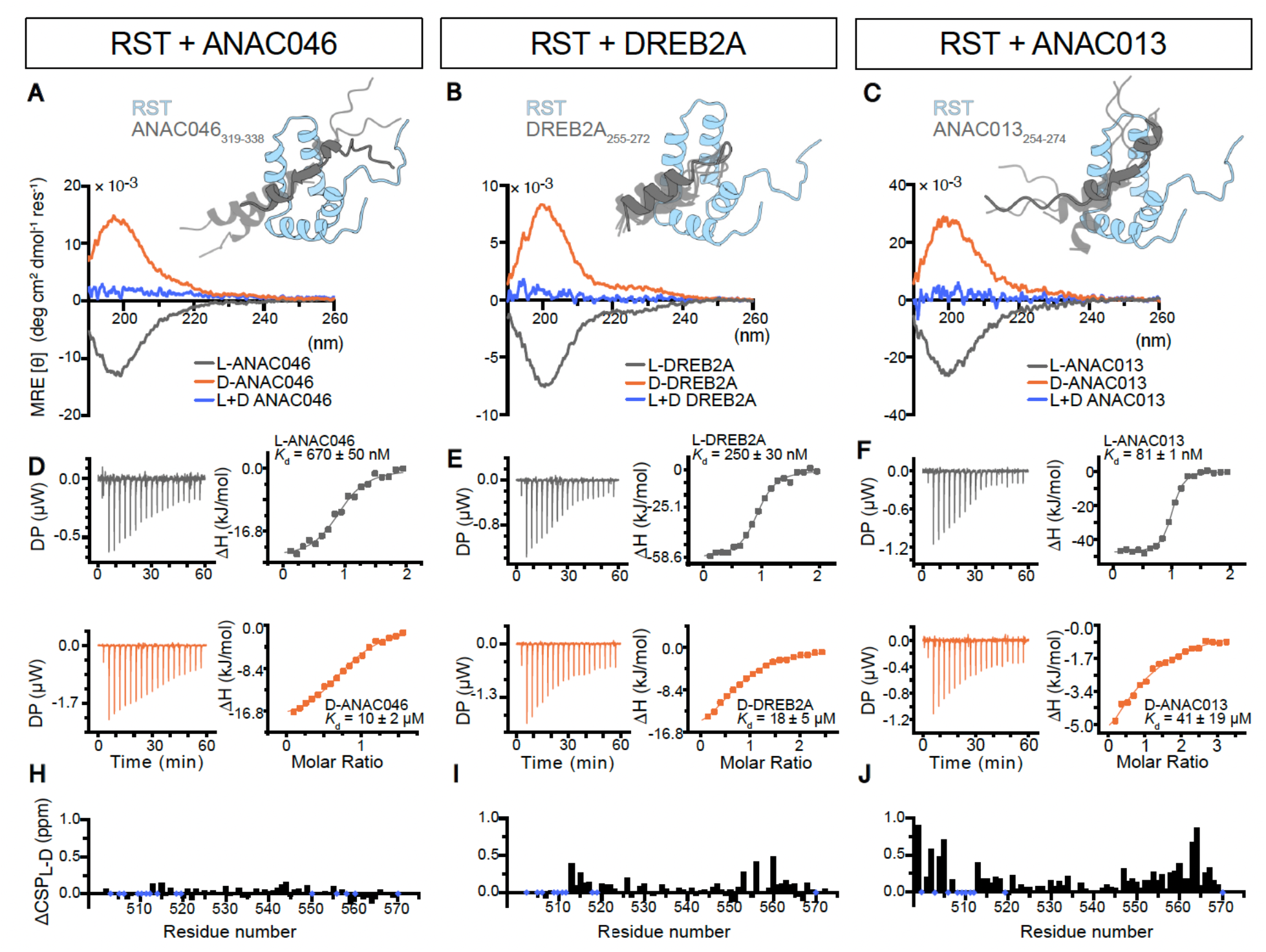
RST interactions with L and D-peptides vary depending on remaining disorder in the complex. **A, B, C** The peptides of ANAC046, DREB2A and ANAC013 are disordered in their free state, and form varying structure upon binding to RST (insets), with far-UV CD spectra showing the L- and D-enantiomers of each peptide as mirror images (blue: sum of L- and D- peptide spectra). **D, E, F** Results obtained from ITC. **H, I, J** NMR CSPs of the interactions showing the differences between the L- and D-enantiomers. Blue diamonds indicate missing assignments, assigned residues that could not be tracked or prolines.

Taken together, a pattern emerges from the data (**Fig. 4**). We find that the differences in average CSPs increase with increasing structure in the bound state (**Fig. 4A**). There is no difference in CSPs between L- and D-H1_155-175_ upon their interactions with ProTα, compared to small differences between the interaction of RST with L- and D-ANAC046 and DREB2A. The CSPs appear to hit a upper limit in the RST:ANAC013 and MCL1:PUMA interactions at an average of ∼0.15 ppm per residue difference between the L- and D-peptides. This upper limit may be due to either reaching a maximum CSP for these proteins or a dependence on the size or properties of the binding site. We also calculated the difference in Δ*G*from ITC between L- and D-peptides (ΔΔ*G*_*D−L*_) for each binding pair (**Fig. 4B**). Here, we again find no significant difference between L- and D-H1_155-175_ interacting with ProTα, while a higher degree of folding in the bound state leads to larger differences in binding free energy between L- and D-peptides. Thus, the energy for folding following encounter complex formation is increased for the D-peptide when the interaction requires more structure. To relate the difference in energy between binding an L- or a D-peptide to the strength of the interaction, we normalized ΔΔG_D-L_ to ΔG_L_ (**Fig. 4C**). This shows again that the energy difference between binding an L- or D-peptide scales with the degree of order and disorder. We also calculated the change in *K*_d_, with no change between the L- and D-H1_155-175_ for ProTα, and with D-ANAC013 having a 500-fold higher *K*_d_ than L-ANAC013 for RST (**Table S3**). In comparing ΔH and TΔS for the RST interactions, we observe a linear relationship between the changes in enthalpy and entropy across the binding partners, with disordered and D-peptide interactions producing smaller changes in enthalpy and entropy (**Fig. 4D**), similar to previous studies comparing folding-upon-binding complexes^29^. When we represent these data using propensity for interaction, ProTα has equal propensity of interacting with L- or D-H1_155-175_, whereas RST and MCL1 only have a 10% propensity of interaction with D-ANAC013 and D-PUMA, respectively (**Fig. 4E**). The difference in propensity most likely comes from non-productive encounters that lead to dissociation and not binding. Free energy schematics of the protein interactions using calculated ΔΔG_(D-L)_ and ΔΔG_unbound-‡,(D-L)_-values illustrate that RST:ANAC013 becomes energetically unfavorable when the ligand is the D-enantiomer (**Fig. 4F**). In summary, it becomes apparent that the propensity for interaction with a D-peptide, and thus independence of stereochemistry, directly relies on the extent of disorder in the complex.

**Figure 4.**
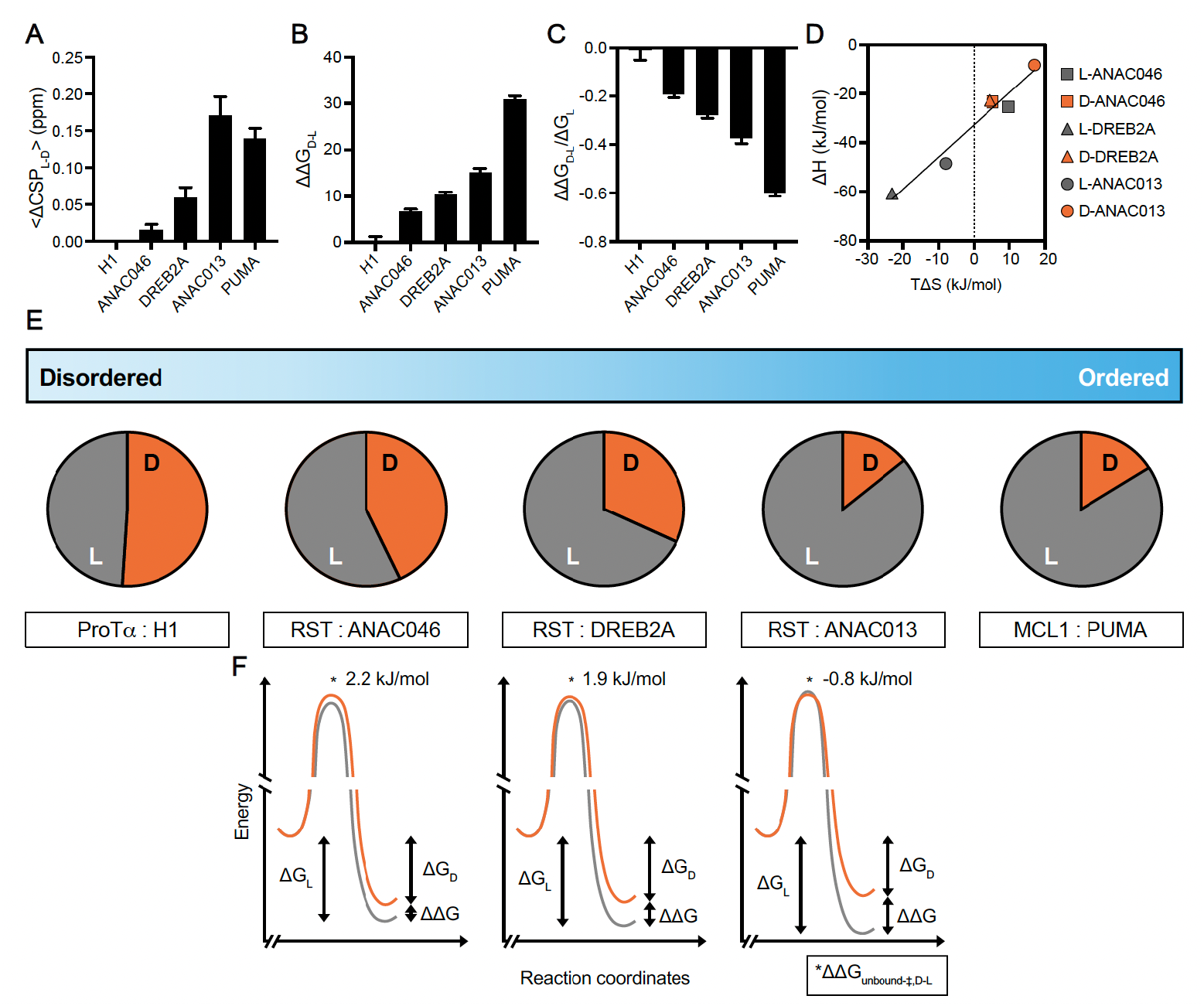
Remaining disorder in complexes scales with the ability to bind D-enantiomers. **A** The difference in average CSPs caused by D- or L-peptides in each system. **B** The difference in ΔG (ΔΔG_D-L_) recorded via ITC for L- or D-peptides in each system. **C** The difference in ΔG (ΔΔG_D-L_) normalized to the strength of the interaction (ΔG_L_). **D** Linear fit of the relationship between entropy (TΔS) and enthalpy (ΔH) for RST interactions. **E** Scale of disordered to ordered protein interaction systems with representative probabilities of each protein interacting with a L- or D-peptide based on differences in CSPs. ProTα has almost equal probability of interacting with D- or L-H1_155-175_, whereas RST and MCL1 have ∼10% probability of interacting with D-ANAC013 or D-PUMA, respectively. **F** Free energy diagrams of transcription factor interactions with RST (orange: D; gray: L) with differences in binding free energies, ΔΔG, from **B** and differences in activation free energies between D- and L-peptides, ΔΔG_unbound-‡,D-L_, from NMR lineshape analysis (**Table S3**). Error bars indicate SEM.

## DISCUSSION

It is counterintuitive that an L-protein should be able to interact with a D-version of its natural partner. However, when considered further, we asked whether and why this applies to a truly disordered interaction. We find that protein complexes that are fully disordered (e.g. ProTα:H1) form regardless of chirality. To our surprise, the observation was not limited to completely disordered polyelectrolyte binding partners. Instead, the propensity for interaction between L- and D-proteins exists on a continuum of disorder, irrespective of charge, hydrophobicity, or other features (**Table S1**). We also found evidence of an encounter complex between D-PUMA and MCL-1, a state which has historically been difficult to observe^30^. Overall, we have identified a method of inferring the degree of disorder within a protein complex by its sensitivity to chirality, applicable not only in the case of a fully disordered complex driven by electrostatics. This finding has translational applications in drug design and implications for our understanding of protein evolution.

Chirality is an important feature for interactions relying on structure or the formation of structure, but an electrostatic, disordered interaction can proceed regardless. Each of the peptides studied here relies on electrostatics to interact with their respective binding partner^16,18,31^, therefore, the determining factor appears to be the degree of disorder. PUMA forms an encounter complex with MCL1 due to long-range electrostatics^31^, which is likely the reason we observe loss of MCL1 peak intensities induced by D-PUMA encounters. Electrostatics are also integral to the interactions of RST^18^. Initially, based on structural predictions^27^ and CD data^26^, we expected more structure to form in the interaction between RST and DREB2A compared to the complex of RST and ANAC013. While our results for ANAC046 supported remaining disorder in the complex, we observed a complex between RST and ANAC013 that was more structured than expected from previously published data^26^. This discrepancy is likely caused by the difficulty in predicting the interactions and dynamics of disordered proteins^32,33^ and from our internal bias towards searching for structure. Our results are therefore also relevant in the context of chaperones where one protein is ordered and the other disordered, as seen in the chaperone (GroEL/ES) which can assist the folding of both L- and D-enantiomers^34^.

In this study, we have investigated the disorder-order continuum of protein interactions with relatively similar electrostatic and hydrophobic features (**Table S1**). However, we do not know whether fully disordered hydrophobic complexes exist and whether they are sensitive to chirality, although hydrophobic ligands have been shown to remain disordered in complexes^35^. Research into highly hydrophobic D-amino acid-based transcriptional coactivators has demonstrated that they can induce transcription to a similar level as their L-enantiomer counterpart^36^, suggesting that at least some hydrophobic disordered proteins can still be functional regardless of chirality, and bind with high affinity. Thus, as IDPs have been historically difficult to target, requiring novel strategies^37,38^, the presented results have substantial implications for the development of new, stable D-peptide drugs directly binding to IDPs.

Protein-protein interactions are fundamental for sustaining the information network that separates life from non-living systems. For simplicity, we assume that the distribution of abiotic amino acids of either chirality in the “primordial soup” was similar^39^. Moreover, peptide bonds may form equally well between L- and D-, D- and D, or L- and L-amino acids^40^. Therefore, peptide-peptide interactions between heterochiral peptides can be envisaged to have existed before biological systems became homochiral^41^. The herein described example of a chirality-independent disordered complex is the first one reported and is an exciting hint at a living entity existing before chiral preferences evolved, before the first replicator arose, and therefore before the Last Universal Common Ancestor.

## Acknowledgements

The authors thank Kristian Strømgaard for valuable discussions on D-peptide synthesis, Christian B. Parsbæk for initial discussions on the MCL1:PUMA complex, Andreas Prestel for expert NMR assistance and Signe S. Sjørup and Charlotte O’Shea for technical support and protein purification. We thank Alexander S. Hebbelstrup, Christian B. Parsbæk, and Kaare Teilum for the MCL1 construct. Julia Diaz is thanked for graphical support and Daniel Nettels for providing data analysis tools. This work was supported by the Novo Nordisk Foundation challenge grant REPIN, rethinking protein interactions (NNF18OC0033926 to K.S., B. S. and B.B.K.), by the Danish Research Councils (9040-00164B to B.B.K.), and by the Swiss National Science Foundation (to B.S.). Further support was given by the European Union’s Horizon 2020 research and innovation programme under the Marie Sklodowska-Curie grant agreement No. 101023654 (awarded to E.A.N.). NMR spectra were recorded at cOpenNMR, an infrastructure facility funded by the Novo Nordisk Foundation (NNF18OC0032996).

## Author contributions

This study was conceptualized by J.G.O., E.A.N., A.D.D., L.S., K.B., K.S., and B.B.K.; J.G.O., L.S., K.B. and C.R.O.B. designed the D-peptides; E.A.N. carried out experiments on ProTα and MCL1 presented in this manuscript, with initial experiments on ProTα conducted by C.B.F. and K.B. smFRET was performed by A.S. in collaboration with B.S.; A.D.D. carried out experiments with RST presented in the manuscript, with initial experiments conducted by E.D., I.B. and L.S.; E.A.N., A.D.D., K.S., J.G.O. and B.B.K. wrote the manuscript with contributions from all authors. All authors have read and agreed to the submitted/published version of the manuscript.

## Additional information

The online version contains supplementary material available at xxx

## Data availability

Chemical shifts of MCL1 in the PUMA bound state have been submitted to BioMagResBank under the accession number 52264.

## METHODS

### Synthetic peptides

Synthetic L- and D-peptides of H1_155-175_, PUMA_130-156_, ANAC013_254-274_, ANAC046_319-338_, and DREB2A_255-272_ were purchased from Pepscan (NL) (now Biosynth, US) at a purity of minimum 95% and purified by HPLC. The D-peptides contain amino acid residues with a stereoisomeric D-form of each chiral carbon. The peptides were either resuspended in MilliQ H_2_O or in MilliQ H_2_O containing 50 mM NH_4_HCO_3_ and lyophilised repeatedly to remove leftover trifluoroacetic acid from the last purification step by the manufacturer. Peptides were then either resuspended directly in the buffer used for experiments or in H_2_O w/o 50 mM NH_4_HCO_3_ to measure the concentration. If no aromatic residue was present in the peptide sequence, the absorbance at 214nm was used. The extinction coefficient was calculated using Bestsel^42^.

### Expression and purification of proteins

^15^N-labelled and unlabelled full-length ProTα was expressed and purified as described^15^. The double-cysteine variant of ProTα (E56C/D110C) used in smFRET experiments was expressed and purified as described^16^, with some modifications. Briefly, ProTα was dialyzed against Tris buffer (50 mM Tris, 200 mM NaCl, 2 mM DTT, 1 mM EDTA; pH 8), during which the hexa-histidine tag was cleaved using HRV 3C protease. Cleaved ProTα was purified further using Ni Sepharose Excel resin (Cytiva, formerly GE Healthcare; Søborg, Denmark) and a HiPrep Q FF column (Cytiva) with a gradient from 200 mM to 1 M NaCl. Buffer was exchanged (HiTrap Desalting column (Cytiva)) to labeling buffer potassium phosphate (100 mM; pH 7). ^15^N-labelled and unlabelled GST-MCL1_152-308_ was expressed in BL21(DE3)pLysS *E. coli* in the presence of ampicillin. Cells were grown at 37 °C in LB or M9 minimal media (for ^15^N-labelling) until OD_600_ reached 0.6, then induced with IPTG (1 mM final concentration) and harvested after four hours. The cell pellet was resuspended in Tris buffer (20 mM Tris, 100 mM NaCl; pH 8), then lysed by sonication. After pelleting again, the supernatant was applied to GST Sepharose beads (Cytiva), and GST-MCL1_152-308_ was eluted using Tris-GSH buffer (20 mM Tris, 100 mM NaCl, 10 mM GSH; pH 8). The GST-tag was removed using TEV protease (0.7 mg) overnight at room temperature. Final purity was reached using a Superdex 75 26/60 column (Cytiva), equilibrated with 50 mM phosphate buffer (pH 7). ^13^C-^15^N-labelled MCL1_152-308_ was expressed as described^43^ and purified as above. The expression and purification of ^15^N-labelled and unlabelled RCD1-RST_499-572_ were carried out as previously described^18^ with the lysis buffer changed to 20 mM Tris-HCl, pH 9.0, 20 mM NaCl. The buffer used in the last purification step by size exclusion chromatography on a Superdex 75 10/300 GL column (Cytiva) was the buffer described for the individual methods.

### Far-UV circular dichroism (CD) spectropolarimetry

Far-UV CD spectra of L- and D-peptides of H1_155-175_, PUMA_130-156_, ANAC013_254-274_, ANAC046_319-338_, and DREB2A_255-272_ were measured on a Jasco 815 spectropolarimeter with a Jasco Peltier control in the range of 260-190 nm at 20 °C. Concentrations of peptides varied between 10-30 μM in either MilliQ H_2_O, pH 7.0 (PUMA_130-156_, H1_155-175_) or 20mM NaH_2_PO_4_/Na_2_HPO_4_, pH 7.0 (ANAC013_254-274_, ANAC046_319-338_, DREB2A_255-272_) with 1 mM TCEP in the samples containing ANAC046 peptides. A quartz cuvette with a 1mm path length was used and 10 scans were recorded and averaged with a scanning speed of 20 nm/min and response time of 2 sec. A spectrum of the buffer using identical setting was recorded for each protein and subtracted the sample spectrum. Spectra were not averaged or smoothed.

### NMR spectroscopy

All NMR spectra were recorded on Bruker Avance III 600 MHz, 750 MHz or 800MHz (for H) spectrometers equipped with cryoprobes. Natural abundance H, ^15^N and H, ^13^C-HSQC spectra were recorded on all peptides at either 10 °C or 25 °C to ensure stereoisomeric properties. Peptides (0.5 mM) in sample buffer containing 20 mM Na_2_HPO_4_/NaH_2_PO_4_ pH 7.0, 100 mM NaCl, 10 % (v/v) D_2_O, 0.02 % (w/v) NaN_3_ and 0.7 mM 4,4-dimethyl-4-silapentane-1-sulfonic acid (DSS) for ANAC046_319-338_, ANAC013_254-274_ and DREB2A_255-272_ with the addition of 1 mM DTT in the samples containing ANAC046 peptides. H, ^15^N-HSQC spectra were recorded on 50 μM ProTα, with or without 500 μM L- or D-H1_155-175_ in TBSK (ionic strength 165 mM; pH 7.4). H, ^15^N-HSQC spectra were recorded on 50 μM MCL1, with or without 100 μM L- or D-PUMA_130-156_, in Tris (50 mM; pH 7.0). Assignments of ^13^C,^15^N-MCL1 interacting with L-PUMA_130-156_ were completed from a series of HNCACB and HNCOCACB 3D spectra as described ^44^, and deposited to BMRB (52264). H, ^15^N-HSQC spectra were recorded on ^15^N-labelled 100 μM RCD1-RST_499-572_ in 20 mM Na_2_HPO_4_/NaH_2_PO_4_ pH 7.0, 100 mM NaCl, 10 % (v/v) D_2_O, 0.02 % (w/v) NaN_3_ and 0.7 mM DSS at 25 °C in the absence and presence of each stereoisomeric forms of 0-200μM ANAC046_319-338_, ANAC013_254-274_ and DREB2A_255-272_ in the following ratios; 1:0, 1:0.2, 1:0.4, 1:0.6, 1:0.8, 1:1 and 1:2. Assignments of ProTα and RCD1-RST were taken from BMRB entries 27215 and 50545, respectively^15,18^.

Amide CSPs were calculated from the H, ^15^N-HSQCs in the absence and presence of the highest concentration of peptide used for each interaction using Equation 1:

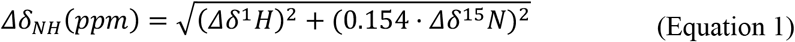

The difference in CSPs (ΔCSPs) induced by either L- or D-peptides was calculated per residue (L-D) and then averaged over the whole protein, not including residues which could not be tracked.

#### 2D NMR lineshape analysis

2D NMR lineshape analyses were performed for interactions of L-and D-peptides with RCD1-RST_499-572_. The recorded H, ^15^N HSQC spectra were processed using qMDD with exponential weighting functions with 4Hz and 8Hz line broadening in the direct and indirect dimensions, respectively. The 2D lineshape analysis was performed using the tool TITAN^45^ in Matlab (Mathworks; Sweden). All titrations were fitted to a two-state binding model, and at least 12 spin systems were picked for each analysis. Due to initial poor fitting for the titrations of the interaction ^15^N-RCD1-RST_499-572_ and L-ANAC013_254-274_, the *K*_d_ was fixed using the values determined from ITC. Errors were determined by a bootstrap analysis using 100 replicas to determine the standard deviation from the mean. From the lineshape analysis, the fitted *K*_d_ and *k*_*off*_ were used to calculate *k*_*on*_ based on equation 2:

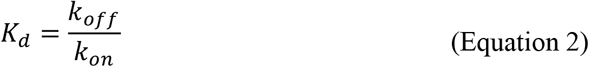

The differences in activation free energies for binding between D- and L-peptides were estimated from the ratios of the association rate constants for both stereoisomers, 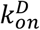and 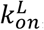, based on Equation 3:

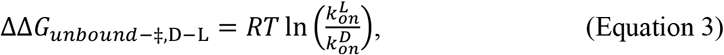

which was rewritten from Fersht (Equation 18.22)^46^.

### Isothermal titration calorimetry (ITC)

Prior to ITC, all samples were spun down at 17,000 rpm for 10 min at the experimental temperature. ITC experiments involving ProTα and MCL1_152-308_ as interaction partners were recorded on MicroCal PEAQ-ITC microcalorimeter (Malvern, Malvern, United Kingdom). ProTα (7.1 μM) was placed in the cell and either L- or D-H1_155-175_ (99.1 μM) in the syringe, in TBSK (165 μM ionic strength) at 20 °C. Each injection was 2 μL, with a total of 19 injections at an interval of 150 s between each. Data were fit using a fixed number of binding sites (fixed to one) so that fits could be standardized. For the MCL1_152-308_ interactions, MCL1_152-308_ (10 μM) was placed in the cell, with either L- or D-PUMA_130-156_ (100 μM) in the syringe, in Tris (50 mM; pH 7.0) at 25 °C. Each of the 35 injections was 1 μL, with an interval of 150 s between each. The experiment was repeated for MCL1:D-PUMA_130-156_, increasing the concentrations to 70 and 700 μM, respectively, while keeping the remaining experimental conditions identical. ITC experiments involving RCD1-RST_499-572_ as interaction partner were recorded on a MicroCal™ ITC_200_ microcalorimeter (Cytiva) at 25 °C in 50mM Na_2_HPO_4_/NaH_2_PO_4_ pH 7.0, 100mM NaCl. 1 mM TCEP was added the sample buffer for interactions involving ANAC046 peptides. Concentrations of RCD1-RST (499-572) varied between 10-100 μM in the cell and 100-1000 μM of the ANAC046, ANAC013 or DREB2A peptides in the syringe. The first injection was 0.5 μL followed by 18 repetitions of 2 μL injections separated by 180 seconds. These experiments were processed using the Origin7 software package supplied by the manufacturer. The last 18 injections of each experiment were fitted to a one set of sites binding model. Triplicates were recorded for each interaction.

### Fluorophore labelling for smFRET

ProTα was labelled by incubating it with Alexa Fluor 488 (0.7:1 dye to protein molar ratio) for 1 hour at room temperature and sequentially with Alexa Fluor 594 (1.5:1 dye to protein molar ratio) overnight at 4 °C. Labelled protein was purified using a HiTrap Desalting column and reversed-phase high-performance liquid chromatography (RP-HPLC) on a SunFire C18 column (Waters Corporation, Baden-Dättwil, Switzerland) with an elution gradient from 20% acetonitrile and 0.1% trifluoroacetic acid in aqueous solution to 37% acetonitrile. ProTα-containing fractions were lyophilized and dissolved in buffer (10 mM Tris, 200 mM KCl, 1 mM EDTA; pH 7.4).

### Single-molecule FRET measurements and analysis

Single-molecule fluorescence experiments were conducted using either a custom-built confocal microscope or a MicroTime 200 confocal microscope (PicoQuant, Berlin, Germany) equipped with a 485-nm diode laser and an Olympus UplanApo 60x/1.20 W objective. Microscope and filter setup was as previously described^16^. The 485-nm diode laser was set to an average power of 100 μW (measured at the back aperture of the objective), either in continuous-wave or pulsed mode with alternating excitation of the dyes, achieved using pulsed interleaved excitation (PIE). The wavelength range used for acceptor excitation in PIE mode was selected with a z582/15 band pass filter (Chroma, Olching, Germany) from the emission of a supercontinuum laser (EXW-12 SuperK Extreme, NKT Photonics, Landsberg am Lech, Germany) driven at 20 MHz, which triggers interleaved pulses from the 485-nm diode laser used for donor excitation. In our experiments, photon bursts (at least 3000 bursts) were selected against the background mean fluorescence counts and, in case of pulsed interleaved excitation, by having a stoichiometry ratio *S* of 0.2<*S*<0.75, each originating from an individual molecule diffusing through the confocal volume. Transfer efficiencies were quantified according to *E* = *n*_A_⁄(*n*_A_+*n*_*D*_), where *n*_*D*_ and *n*_A_ are the numbers of donor and acceptor photons in each burst, respectively, corrected for background, channel crosstalk, acceptor direct excitation, differences in quantum yields of the dyes, and detection efficiencies. All smFRET experiments were performed in μ-Slide sample chambers (Ibidi, Germany) at 22 °C in TEK buffer with an ionic strength of 165 mM fixed with KCl; 140 mM 2-mercaptoethanol and 0.01% (v/v) Tween-20 were added for photoprotection and for minimizing surface adhesion, respectively. Single-molecule data were analysed using the Mathematica (Wolfram Research) package Fretica (https://schuler.bioc.uzh.ch/programs). For quantifying binding affinities, transfer efficiency histograms were constructed from single-molecule photon bursts identified as described above. Each histogram was normalized to an area of 1 and fit with a Gaussian peak function to extract its mean transfer efficiency ⟨*E*⟩. Consequently, the mean transfer efficiency as a function of increasing concentration of D/L-H1_155-175_, ⟨*E*⟩(*C*_*D*/*L*-H1_), was fit with:

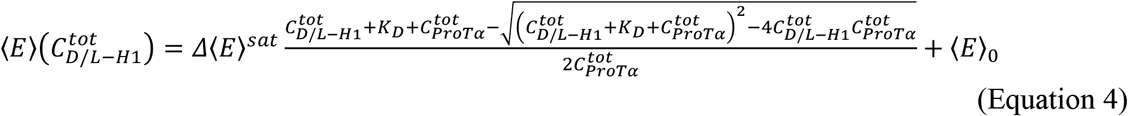

Here, 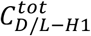 and 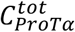 are the total concentration of D/L-H1_155-175_ and ProTα, respectively, ⟨*E*⟩_0_ is the mean transfer efficiency of free ProTα, and Δ⟨*E*⟩^*sat*^ is the increase in transfer efficiency of ProTα saturated with D/L-H1_155-175_, while *K*_*D*_ is the binding dissociation constant.

## SUPPLEMENTARY FIGURES

**Figure S1.**
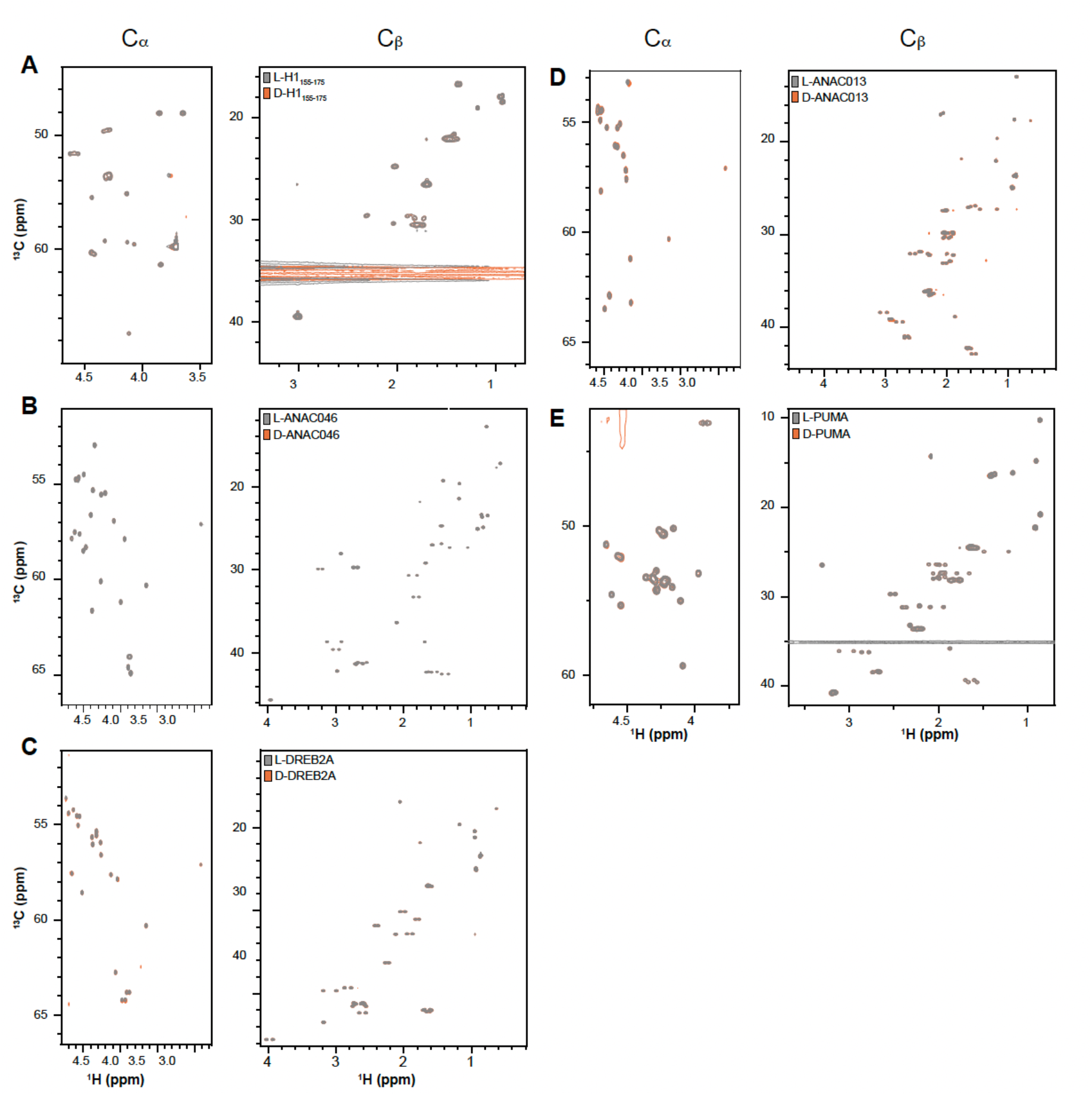
^13^C-HSQC NMR spectra showing Cα and Cβ chemical shifts of L- and D-peptides. **A** L-H1_155-175_ and D-H1_155-175_; **B** L-ANAC046 and D-ANAC046; **C** L-DREB2A and D-DREB2A; **D** L-ANAC013 and D-ANAC013; **E** L-PUMA and D-PUMA. All L-peptides displayed in grey and D-peptides in orange.

**Figure S2.**
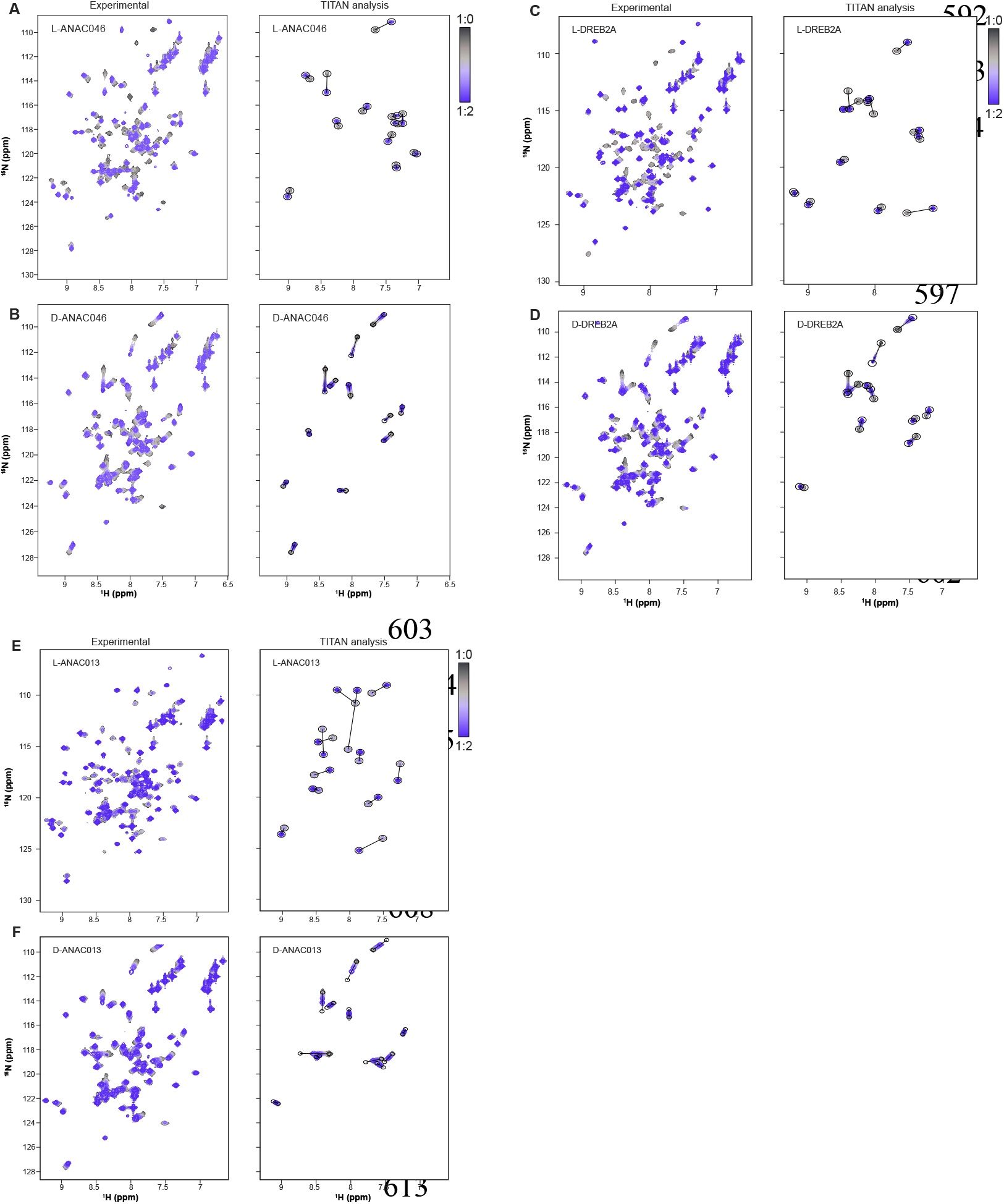
NMR lineshape analysis of titration of RST with RST-interacting peptides using TITAN^45^. **A** L-ANAC046; **B** D-ANAC046; **C** L-DREB2A; **D** D-DREB2A; **E** L-ANAC013; **F** D-ANAC013.

## SUPPLEMENTARY TABLES

**Table S1.**
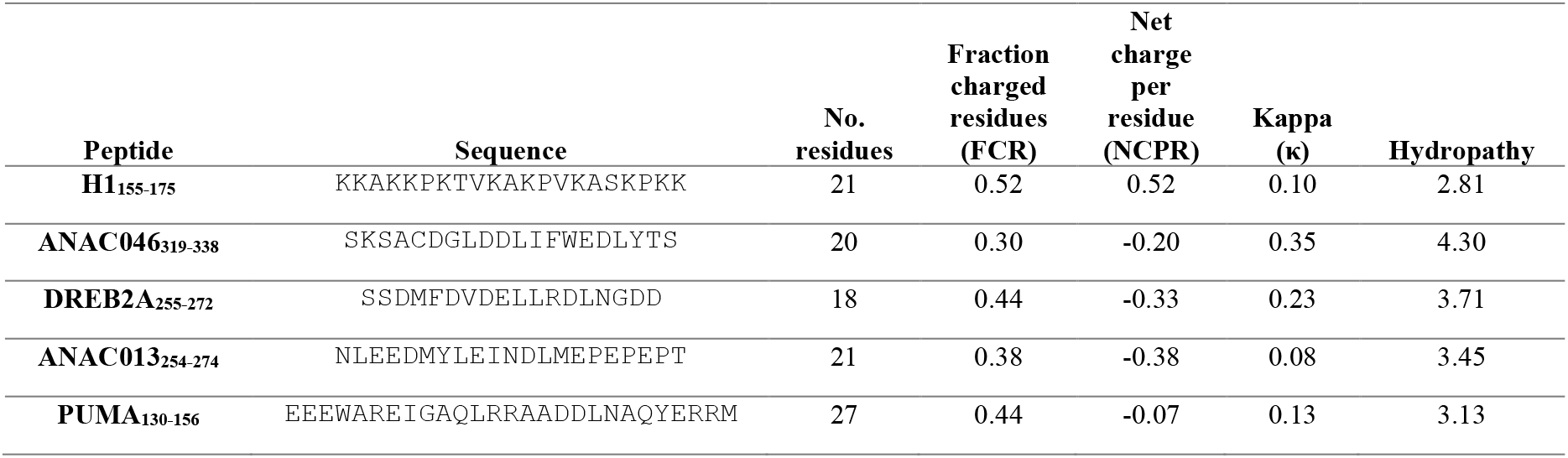
CIDER^47^ analysis of disordered peptides.

**Table S2.**
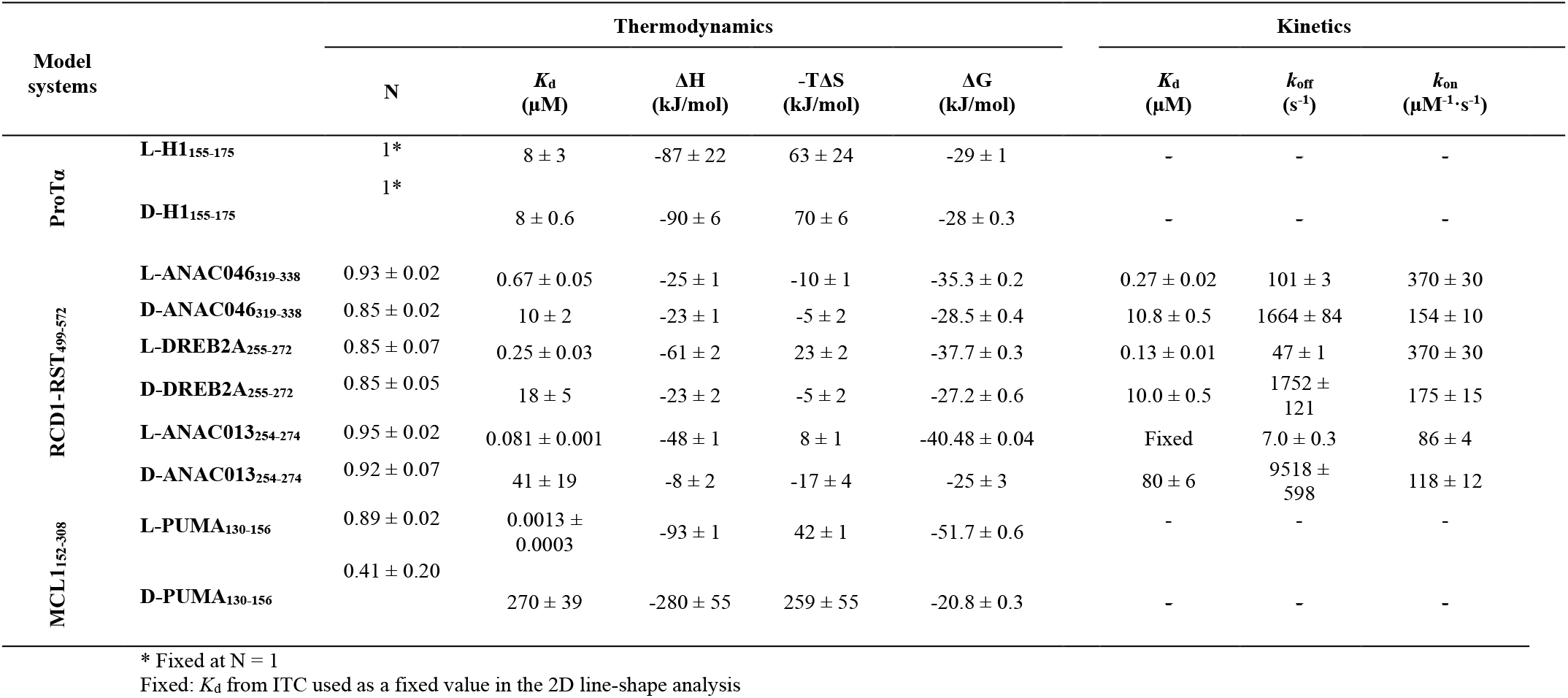
Thermodynamics and kinetics for L- and D-peptide interactions.

**Table S3.**
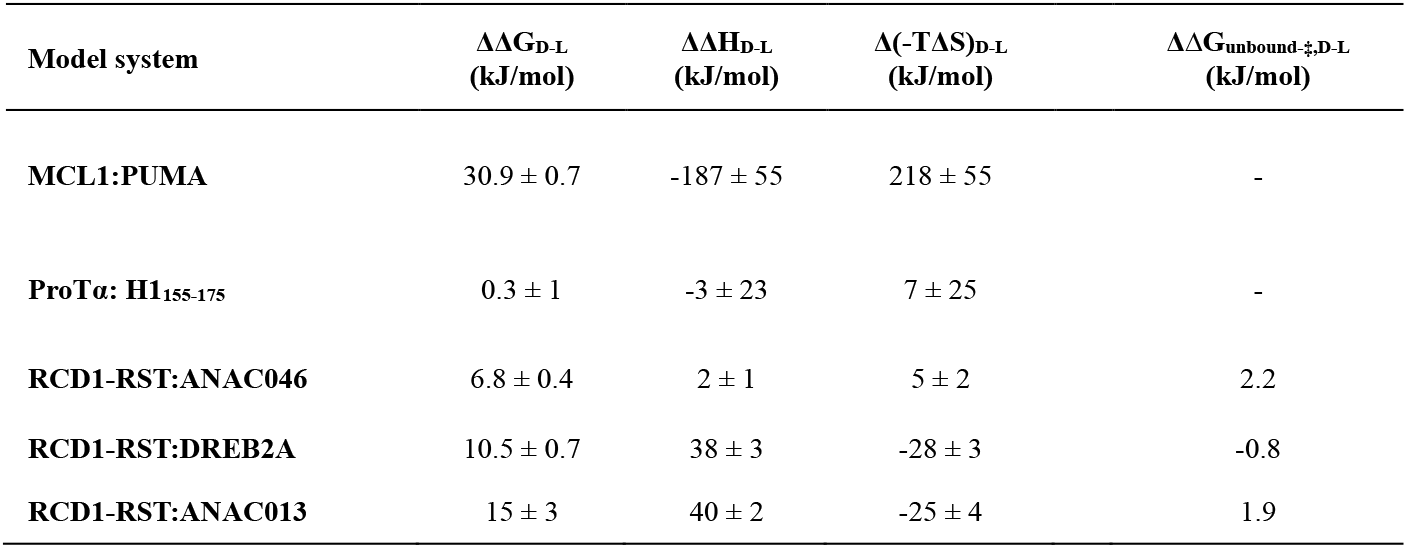
Effects of stereochemistry on interactions.

